# Simulated musculoskeletal optimization for sprinting and marathon running

**DOI:** 10.1101/2023.08.07.552222

**Authors:** Tom Van Wouwe, Jennifer Hicks, Scott Delp, Karen Liu

**Affiliations:** Department of Bioengineering, Stanford University; Department of Computer Science, Stanford University

## Abstract

Musculoskeletal geometry and muscle volumes vary widely in the population and are intricately linked to the performance of tasks ranging from walking and running to jumping and sprinting. However, our ability to understand how these parameters affect task performance has been limited due to the high computational cost of modelling the necessary complexity of the musculoskeletal system and solving the requisite multi-dimensional optimization problem. For example, sprinting and running are fundamental to many forms of sport, but past research on the relationships between musculoskeletal geometry, muscle volumes, and running performance has been limited to observational studies, which have not established cause-effect relationships, and simulation studies with simplified representations of musculoskeletal geometry. In this study, we developed a novel musculoskeletal simulator that is differentiable with respect to musculoskeletal geometry and muscle volumes. This simulator enabled us to find the optimal body segment dimensions and optimal distribution of added muscle volume for sprinting and marathon running. Our simulation results replicate experimental observations, such as increased muscle mass in sprinters, as well a mass in the lower end of the healthy BMI range and a higher leg-length-to-height ratio in marathon runners. The simulations also reveal new relationships, for example showing that hip musculature is vital to both sprinting and marathon running. We found hip flexor and extensor moment arms were maximized to optimize sprint and marathon running performance, and hip muscles the main target when we simulated strength training for sprinters. Our simulation results can help sprint and marathon runners customize strength training, and our simulator can be extended to other athletic tasks, such as jumping, or to non-athletic applications, such as designing interventions to improve mobility in older adults or individuals with movement disorders.

**AUTHOR SUMMARY:** Our study addresses the challenge of determining optimal musculoskeletal parameters for tasks like sprinting and marathon running. Existing research has been limited to observational studies and simplified simulations. To overcome these limitations, we developed a differentiable musculoskeletal simulator to optimize running performance. We replicated past findings and uncovered new insights. We confirmed the benefits of increased muscle mass for sprinters and identified key factors for marathon runners, such a mass in the lower end of the healthy BMI range and an increased leg-length-to-height ratio. Hip musculature was found to be critical for both sprinting and marathon running.

Our simulation results have practical implications. They can inform customized strength training for sprinters and marathon runners. Additionally, the simulator can be extended to other athletic tasks, benefiting various sporting events. Beyond athletics, our open-source simulator has broader applications. It can determine minimal strength requirements for daily activities, guide strength training in the elderly, and estimate the effects of simulated musculoskeletal surgery.

## INTRODUCTION

Performance of tasks ranging from rising from a chair to competing in Olympic-level sporting events depends on precise coordination of many muscles. Variations in an individual’s musculoskeletal geometry and muscle volumes affect performance for many movement tasks [1–3], but our ability to identify cause-effect relationships has been limited because quantifying the effects of changing a person’s musculoskeletal geometry or muscle volumes on task performance is complex.

Musculoskeletal simulation could allow researchers to quantify the effects of variations in body segment dimensions and muscle properties on performance. Simulation allows us to observe the influence of variables that cannot be changed in an experiment (e.g., the height of a runner) and enables us to examine whether specific muscular and skeletal features that have been associated with the performance of a task causally contribute to task performance. Previous simulation studies have opened the door to this possibility but have been limited by simplified representations of the musculoskeletal system [4–6]. The majority use gradient-based trajectory optimization, which requires gradient computations that are computationally expensive because there are typically non-differentiable expressions requiring the use of finite differentiation [7]. To understand the effects of body segment dimensions and muscle properties on task performance using finite differentiation to compute gradients will lead to days or weeks of computation time. Falisse et al. [8] enabled automatic differentiation, a faster alternative to finite differentiation, for musculoskeletal simulation, by replacing non-differentiable expressions by differentiable alternatives. However, the algorithm requires body segment dimensions and muscle properties to be fixed. Our simulator can optimize both body segment dimensions and muscle volumes of a complex, three-dimensional muscle-driven musculoskeletal model for a range tasks. By implementing a musculoskeletal simulator that is fully differentiable with respect to both body segment dimensions and muscle volumes, we have enabled simulations to be completed within hours on a standard computer. To exercise this new simulator, we used it to study the role of body segment dimensions and muscle volumes in sprinting and marathon running performance.

The 100m dash is arguably the most prestigious event in track-and-field [9,10], and marathon running serves as a frontier of human endurance [11,12]. Success in these two events has been associated with different skeletal and muscular features, largely through observational studies. For example, Sedeaud et al. [1] found that mean height, mean body mass index (BMI), and variability in BMI decreased with increasing distance of the event in which male runners specialized. Size, proportions, and other aspects of musculoskeletal geometry have also been associated with an advantage in sports such as speed skating and swimming [2], and cycling [3].

Compared to the general population, sprinters exhibit specialization in musculoskeletal geometry, such as a more limited height range (i.e., sprinters are typically not very short or tall) [13]. Although extremes of height are rare in sprinters [13], body height was not associated with performance in 100m personal best times in a group of sprinters [14]. Body proportions, however, might be important for sprint performance. For example, Tomita et al. [15] found that a higher tibia-to-femur length was associated with better performance in 400m runners. They posit that a higher tibia-to-femur length reduces the leg’s moment of inertia with respect to the hip and thus reduces positive work done by the hip flexors during the swing phase of running. Top sprinters have highly developed musculature for both the lower and upper body, as established via both medical imaging [16] and external measurements [14,17–19]. In a group of competitive sprinters, higher thigh girth, hip strength, and body mass index (BMI) was associated with greater sprint performance as assessed via personal best times [14,20]. Further, higher relative muscle volume in the hip muscles has been shown to differentiate elite sprinters, sub-elite sprinters, and non-sprinters [16]. With hip musculature suggested as a limiter of sprinting performance, stronger hip muscles and a deeper pelvis, which increases the moment arms of hip muscles, might be advantageous.

Compared to the general population, distance runners are shorter, have lower BMIs [17,21] and have a higher tibia-to-femur length ratio [22,23]. In trained distance runners, a higher relative tibia-to-femur length ratio is also associated with better running performance [23], as is a higher relative lower limb length [22,24]. Distance runners, compared to sprinters, have lower maximal isometric knee flexor and extensor torques, in absolute terms and when normalized by body weight [25].

While the observational studies described above reveal associations between performance and musculoskeletal features, they are limited in their ability to identify cause-effect relationships. To overcome this, Deane et al. [26] conducted an interventional experiment, which revealed that hip flexor training significantly improves 40-yd dash times. However, such interventional studies are rare, costly, and difficult to control (e.g., hip flexor training might increase strength of additional muscle groups).

Past simulation studies have explored causal relationships between a musculoskeletal model’s capacity (e.g., maximal joint torques or muscle force) and performance in sprinting [27], jumping [4,5] and gymnastics [28]. However, these studies have been limited in their ability to capture the complexity of the musculoskeletal system as it applies to athletic performance [29]. For example, previous simulation studies that investigated jumping performance did not include muscles [5] or included only six muscles and were restricted to two dimensions [4]. Two simulation studies have revealed how muscle force-length-velocity relationships influence sprinting capacity, but were limited to 2D models driven by nine muscles [30,31]. No simulation studies have comprehensively investigated how differences in body-segment dimensions affect running performance.

In this study, we developed a three-dimensional musculoskeletal simulator that is fully differentiable with respect to both body-segment dimensions and muscle properties to analyze the effects of body-segment dimensions, muscle volume, and distribution of muscle volume on sprinting and marathon running performance. We sought to determine how body-segment dimensions affect maximal sprinting speed and the metabolic cost of running a marathon at moderate speed. We also simulated how optimized, targeted strength training improves sprinting and marathon running performance.

Understanding the influence of body segment dimensions and muscle volume distribution on performance can inform the selection of sports in which to compete and personalize training to help maximize performance. Our simulator is open source and enables researchers to conduct additional studies investigating the relationships between musculoskeletal parameters and increased performance, as in sport, or reduced performance, as can result from injuries, diseases, and disorders.

## RESULTS

We performed predictive simulations of a running gait using a three-dimensional musculoskeletal model with 31 degrees-of-freedom, 92 lower-limb muscles, and 8 torque-actuators for the upper body [32]. The generic model represents a male subject with a stature of 1.81m and mass of 75kg, corresponding to the 73^rd^ and 63^rd^ percentiles for 30 year old males in the United States [33], and that matches the strength of a young healthy subject capable to performing athletic activities. The model has been tested and used in many previous studies, including studies of walking [8], running [8,32] and sprinting [34,35].

The optimization of body segment dimensions and muscle volumes for performance was enabled by adapting the musculoskeletal simulator developed by Falisse and colleagues [36]. First, we implemented a smooth and differentiable version of the OpenSim [37] muscle wrapping algorithm by using a neural network approximation (see Methods for details). Second, we formulated the rigid body dynamics, which govern the movements of the skeleton, to be differentiable with respect to the dimensions, masses, and inertias of the body segments. Simulations in this study took 30 minutes to 4 hours to converge, when running on a laptop (11th Gen Intel Core i9 2.5GHz CPU). Prior to these adaptations, an optimization of the type used in this study took several days. The range of computation times across several similar optimization problems is expected due to the non-convexity and high non-linearity of the optimization problems [38] that stem from muscle-tendon dynamics [39] and contact dynamics [40,41].

To simulate marathon running, we imposed a running speed of 3.33m/s and minimized lower limb energy expended over the marathon distance [8]. To simulate sprinting, running speed was maximized. In all simulations, we optimized the excitations of the muscles and torque actuators to achieve the performance criterion of interest (i.e., marathon or sprint performance). Simulated kinematics and ground reaction forces for running at 3.33m/s and sprinting with the generic model were similar to experimental values, giving confidence in the validity of the musculoskeletal model to address our study aims (see Supplementary Materials Figures S.1, S.2 for marathon running and Figures S.3, S.4 for sprinting). To answer our research questions, we performed simulations that optimized body-segment dimensions (i.e., the three-dimensional dimensions of each segment), or the distribution of added muscle volume (i.e., optimal distribution of muscle volume when adding 5% of the total muscle volume). When scaling the segment dimensions, we increased the mass (and inertia) of that segment assuming constant density, and then changed muscle volume evenly across muscles to be proportional to the overall change in model mass.

### Optimizing model body-segment dimensions improves sprinting (17%) and marathon performance (36%)

Optimizing the model’s body-segment dimensions for sprinting increased maximum speed by 17% (Figure 1A; 9.49 m/s for the sprint-optimized body-segment dimensions compared to 8.13 m/s the generic model). Optimizing body-segment dimensions for marathon running reduced the lower limb energy cost of running a marathon at 3.3 m/s by 36% (Figure 1B; 2074 kcal for the marathon-optimized body-segment dimensions compared to 3267 kcal for the generic model). The sprint-optimized model was heavier than the generic model, but height did not change markedly (Figure 1C, D; 83.6 kg, 1.81 m for the sprint-optimized model compared to 75.2 kg, 1.81 m for the generic model), while the marathon-optimized model was lighter and shorter than the generic model (Figure 1C,D; 54.2 kg, 1.76 m for the marathon-optimized model). Predictions for mass and height fell within the range of values found in the top seven fastest all-time male 100m sprinters and marathon runners (Figure 1C, D).

**Figure 1.**
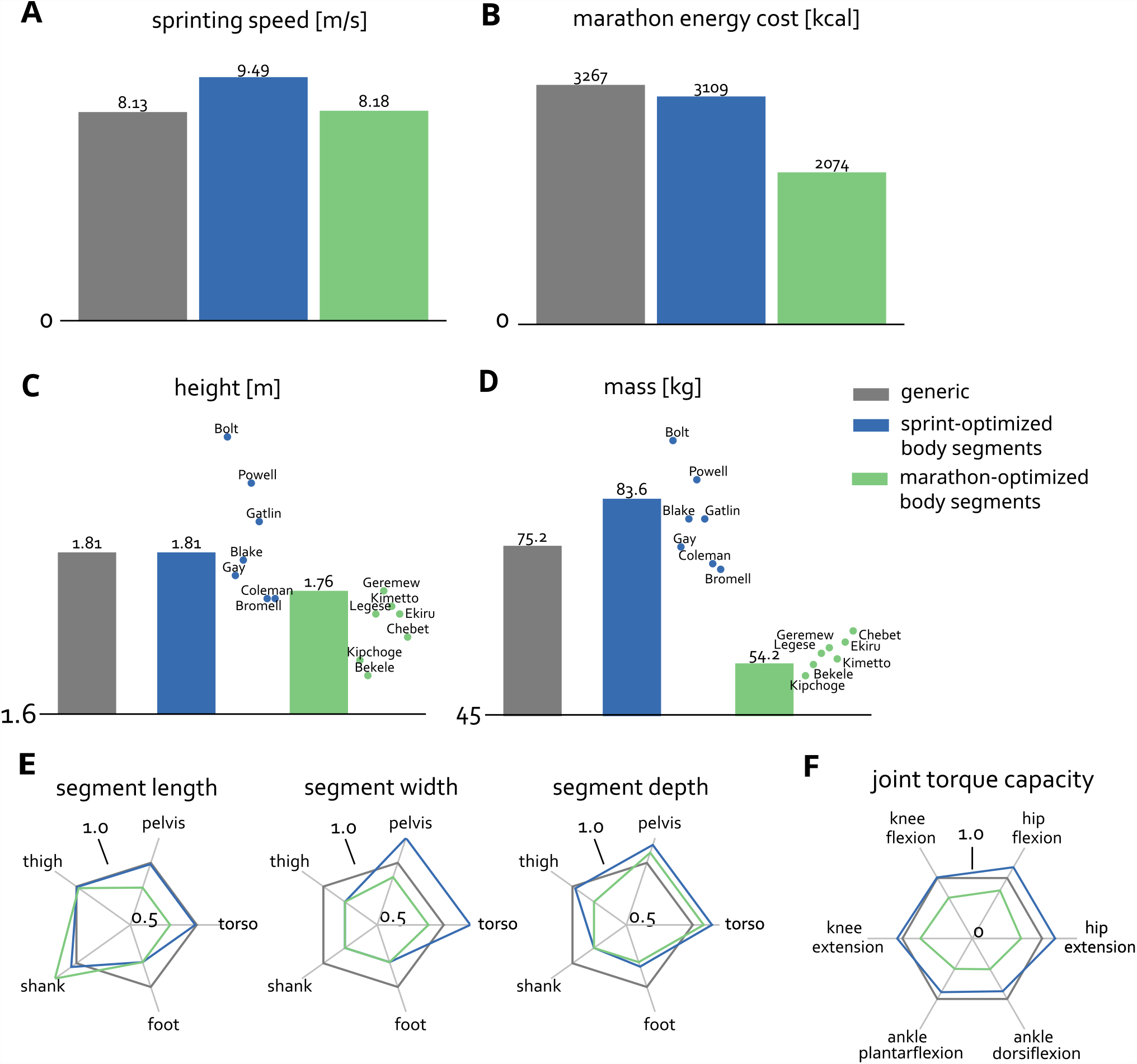
Sprinting speed (A), marathon lower limb energy cost (B), height (C), mass (D) for the generic model and models with optimal body-segment dimensions for sprinting and marathon running. The dots and names in (C) and (D) represent the seven all-time fastest male 100m and marathon runners. In (E) the scaling of individual body segments are shown for the three dimensions: length, width, and depth. The scaling factors are normalized to the generic model. In (F) the figure shows the joint torque capacity of the sagittal plane degrees of freedom for the different models normalized by the capacity of the generic model.

Analysis of the optimized body-segment dimensions and resulting joint-torque capacities revealed the hip muscles as important drivers of sprinting and marathon running performance (Figure 1E,F). The increased pelvis width and depth in both the sprint-optimized and marathon-optimized models (Figure 1E) increased the moment arms of the hip flexors and extensors, leading to increased hip flexion and extension joint torque capacity (Figure 1F). In the sprint-optimized model, the stronger muscles, due to increased mass, further contributed to increased joint-torque capacity. Despite the smaller and weaker muscles in the marathon-optimized model, which were reduced to between 55-75% of the generic model because the model was shorter and thinner, the hip flexion and extension capacities were maintained at about 80% and 72%, respectively, of the generic model due to increased moment arms. At the knee and ankle, the sprint-optimized model showed little change in torque capacity, while the marathon-optimized model showed reduced capacity at both joints, due to lower mass and smaller muscles.

A longer shank was beneficial for both sprinting (+4%) and marathon running (+20%), whereas thigh lengths were maintained in the sprint-optimized model and slightly shorter (1%) in the marathon-optimized model compared to the generic model (Figure 1E). Width and depth of the shank, thigh and foot segments were reduced in both the sprint- and marathon-optimized models, reducing lower-limb inertia.

The sprint-optimized and generic models sprinted with similar step lengths (1.92 m and 1.93 m, respectively), but the sprint-optimized model sprinted with higher step frequency (4.9 Hz vs 4.2 Hz). The marathon-optimized and generic models both ran at marathon pace with a 0.94 m step length and step frequency of 3.5Hz.

### Training the hip muscles and plantarflexors is beneficial for sprinting

We explored how to optimally strengthen muscles by allowing our optimization framework to increase the generic model’s muscle volume with a budget of 5% of the total muscle volume, optimizing for either sprint or marathon performance. We limited the amount an individual muscle’s volume could increase to 20%. The segment dimensions were not changed from the generic model in this analysis.

The model “trained” to optimize sprint performance realized a 9.2% higher sprinting speed (8.88 m/s) compared to the generic model (8.13 m/s) (Figure 2A), while the model “trained” for the marathon reduced the energetic cost of running a marathon by 1.5% compared to the generic model (Figure 2B). Optimized strength training for marathon running was also beneficial for sprinting, evidenced by a 2.3% increase in sprinting speed for marathon-optimized strength. Marathon performance after optimal sprint strength training changed minimally (0.3%). The sprint-optimized model sprinted with a longer step length than the generic model (2.07 m vs. 1.94 m) and a slightly higher step frequency (4.3 Hz vs. 4.2 Hz). The marathon-optimized and generic model both ran at marathon pace with a 0.94 m step length and step frequency of 3.5Hz.

**Figure 2.**
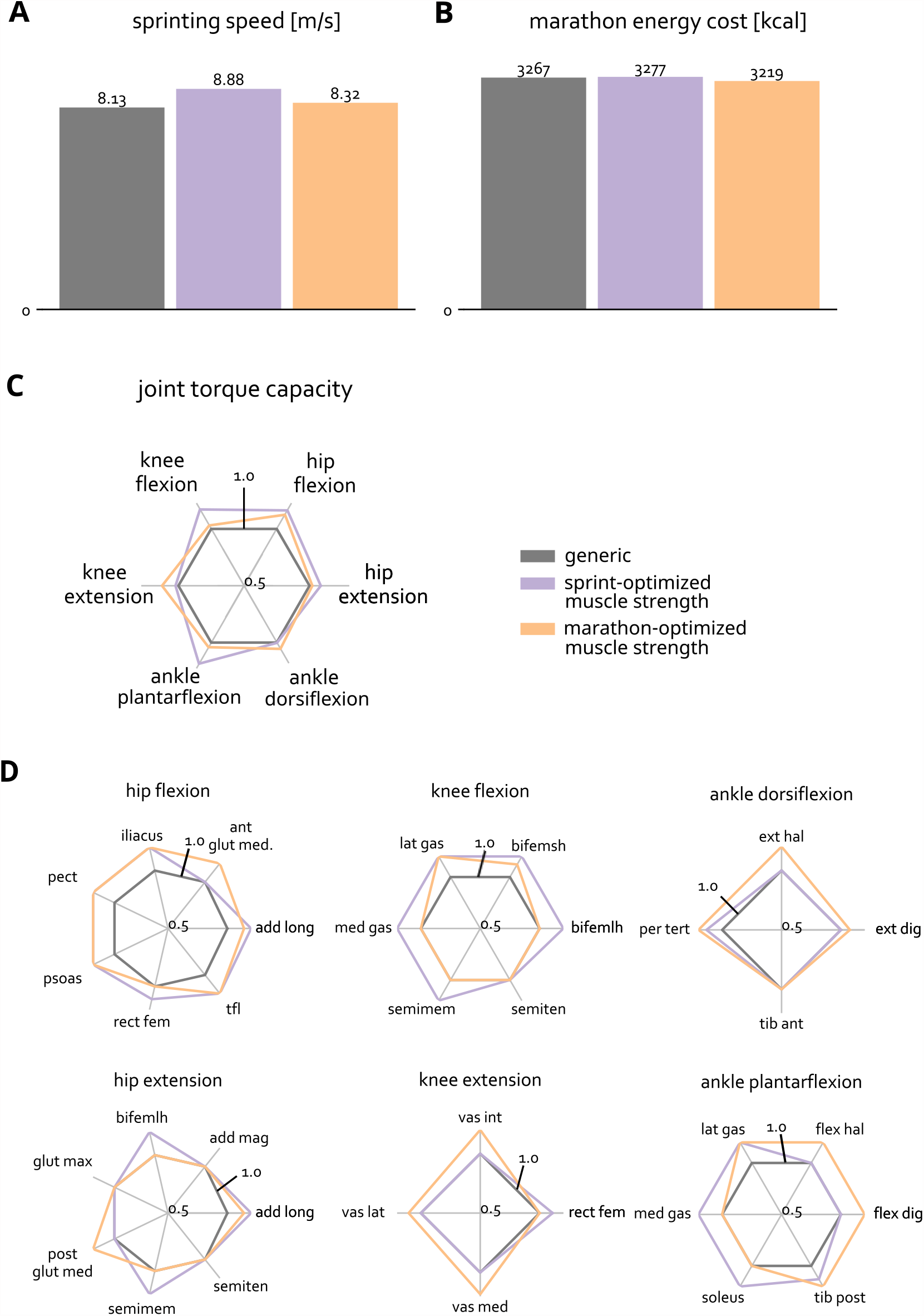
Sprinting speed (A) and marathon lower limb energy cost (B) for the generic model and models after optimal ‘strength training’ for sprinting and marathon running. (C) shows the joint torque capacity of the sagittal degrees of freedom for the different models normalized by the capacity of the generic model. (D) shows normalized muscle maximal isometric force for the different models organized per degree of freedom.

The model with sprint-optimized muscle volumes had increased joint torque capacity primarily for hip flexion, hip extension, knee flexion, and ankle plantarflexion (Figure 2C). Among the muscles with hip flexion moment arms, iliacus, adductor longus, psoas, and tensor fascia latae were beneficial to strengthen (Figure 2D, hip flexion). Among the muscles with hip extension moment arms, biceps femoris longhead, adductor longus, and semimembranosus (Figure 2D, hip extension) were selected as muscles to strengthen. For the ankle plantarflexors, the optimizer suggested strengthening the soleus, and medial and lateral gastrocnemius (Figure 2D, ankle plantarflexion).

We analyzed joint torques during sprinting to understand how the optimized models capitalized on increased muscle volumes. The temporal profiles of the joint torques for the generic model and the two optimized models are similar (Figure 3). However, the model with added muscle volume optimized for sprinting produced a higher peak ankle plantarflexion torque during stance (Figure 3, ankle torque) and a higher peak knee extension torque (Figure 3, knee torque) compared to the generic model. Shortly after take-off, the two optimized models generated more hip flexion torque (Figure 3, hip torque). In terminal swing phase, the sprint optimized model generated more hip extension torque and both optimized models generated higher knee flexion torque (Figure 3, hip and knee torque).

**Figure 3.**
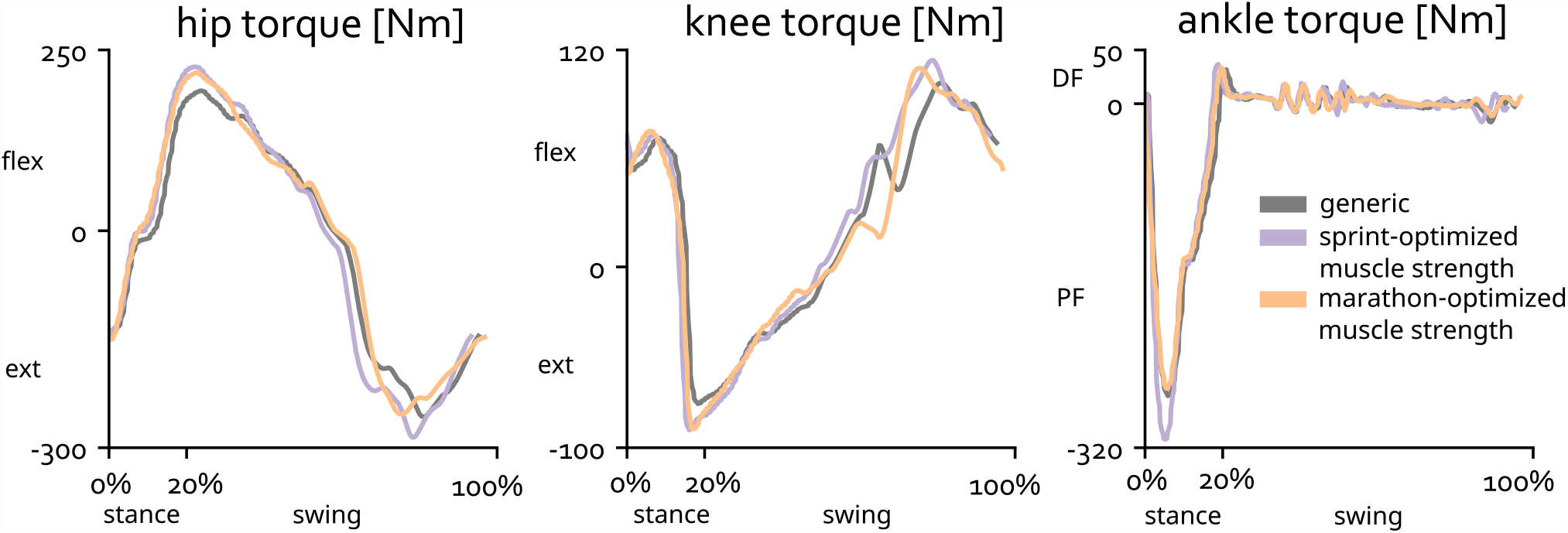
Joint torques for the sagittal-plane degrees of freedom across the sprinting cycle for the right leg. FLEX indicates flexion moment, EXT indicates extension moment, DF indicates dorsiflexion moment, PF indicates plantarflexion moment

## DISCUSSION

Our simulations revealed skeletal geometries and muscle volume distributions that are favorable for sprinting and marathon running, as well as new cause-effect relationships between skeletal geometry, strength, sprinting speed, and marathon running economy. In agreement with experiments [1,16,17,21], sprinting speed improved with additional muscle volume, mainly concentrated at the hip, and marathon economy improved with smaller and lighter body segments and an increased leg-length-to-height ratio. Our simulations also revealed potential mechanisms underlying these relationships. For example, we found that increased hip muscle volume allowed for a faster swing-leg recovery during sprinting, enabling higher step frequency and longer stride lengths. This was the case in our optimizations for both body segment dimensions and muscle volumes. Increased ankle plantarflexor strength was required to generate higher ground reaction forces at these faster sprinting speeds. We also found that an increased shank-to-thigh length ratio and a longer total leg length, without concomitant changes in step length, improved marathon running economy.

Our musculoskeletal simulator is novel since it is differentiable with respect to body-segment dimensions and the inertial properties of a model. We achieved this by (1) formulating the skeleton dynamics to be differentiable with respect to the geometries and inertial properties of the bodies and (2) approximating the computation of muscle wrapping with neural networks, using the musculoskeletal geometry of OpenSim [37] to train the networks. These adaptations allowed us to generate three-dimensional simulations with detailed musculoskeletal models in hours instead of days.

### Musculoskeletal simulation reveals determinants of sprinting performance

Our simulations showed that increasing strength increases sprinting performance. In the case of the model with segment dimensions optimized for sprinting, the increase in performance stemmed from both increases in muscle volume (strength) and optimized geometry. This supports and explains the observation that sprinters, and athletes in sports where sprinting gives an important advantage, are muscular and have an increased BMI [14,16,20].

In support of past correlations observed in experimental and simulation studies [16,34], we identified hip musculature as an important limitation for top sprinting speed. Our sprint-optimized models increased hip flexion and hip extension torque capacities by increasing the moment arms and volumes for these muscles. Our simulated sprinting torques show that faster sprinting goes along with higher hip flexion torques during early swing, to bring the swing leg forward, and higher hip extension torques at the end of swing, to prepare for foot contact. Dorn et al. [34] showed that the hip muscles, primarily the gluteus maximus and hamstrings, are key contributors to propulsion at higher running speeds; we build on this work by showing that increasing the moment-generating capacity of these muscles either by increasing moment arms or increasing muscle volumes indeed leads to faster sprinting speeds.

In alignment with an observational study, which showed that knee flexor strength allowed differentiation between untrained age-matched controls, sub-elite, and elite sprinters [16], knee flexion strength was higher in the sprint-optimized models, with concomitant increases in peak knee flexion torques during sprinting. The increased knee flexor capacity may be a byproduct of the optimizer aiming to maximize ankle plantarflexion capacity, thereby increasing the capacity of the gastrocnemii. Our simulations indicate that knee extension strength may not be a key limiter of sprinting performance, which might explain why the study by Miller et al. [16] could not differentiate elite and sub-elite sprinters based on knee extensor muscle volumes. On the contrary, several other experimental studies associate greater maximal knee extensor torques with faster sprinting [42,43]. However, this does not mean increasing knee extensor strength is required to run faster. In our simulations, both models optimized for sprinting produced greater knee extension torques at midstance, but they did not require a significant increase in knee extension joint torque capacity to do so.

Our sprint-optimized simulations showed that increasing ankle plantarflexion capacity improved sprinting performance, but prior experimental studies are conflicting. Several studies could not correlate ankle muscle volume to sprinting performance [16], [44], but others associated increased ankle plantarflexion strength with increased sprinting performance [27,42]. We found that the model with marathon-optimized body segments slightly increased maximum sprinting speed while having reduced absolute plantarflexion muscle volume, which may help explain the conflicting results in prior experiments.

In agreement with a study by Tomita et al., [15] we found that a long shank and high shank-to-thigh ratio was optimal for sprinting. Because the shank is lighter than the thigh, such geometry decreases lower limb inertia with respect to the hip, increasing the hip flexion acceleration for a given level of hip flexion torque during early swing, without sacrificing leg length. This advantage affects sprint technique, with the faster sprint-optimized model showing increased step frequency rather than longer step length.

### Lower body mass and specialized skeletal geometry affects marathon performance more than increasing strength

When optimizing skeletal geometries, we constrained the body mass index (BMI) to be within a healthy range (between 17.5 and 25.5). The lower limit captures the fact that athletes, and nearly all humans, should maintain a BMI above 17.5 to prevent low bone density [45,46], hormonal issues [47], low energy availability [48], and an array of other health risks [49]. Because our musculoskeletal model and optimization framework does not capture the detrimental performance and health effects of these phenomena, we introduced this lower limit. It is important to note that the optimization did not converge to a minimal achievable mass and predicted a height and mass similar to top marathon runners [1,21]. Mass could have been further reduced within the imposed BMI constraint by lowering shank and thigh length, which would have diminished performance.

Our simulations predicted longer legs, high leg-length-to-height ratio and high shank-to-thigh length ratio as beneficial for marathon running, in line with experimental findings [22,24]. Despite having longer legs, the model with segment dimensions optimized for marathon running maintained stride length compared to the generic model when running at our prescribed marathon pace. This seems counterintuitive, but experimental research has shown that optimal stride length does not correlate well with leg length [50] and is subject specific [51]. Further, stride length varies with speed, requiring different propulsive forces [52]. Reduced muscle strength might explain a smaller optimal stride-to-leg-length ratio for the marathon-optimized model compared to the generic model, since larger steps lead to increased peak joint torques and muscle forces.

The small effects of strength training on marathon performance indicate that strength training is less important for marathon runners than for sprinters. Still, the marathon-optimized body segment dimensions indicate hip musculature as a potential limiter, as hip flexion and extension capacity was mostly maintained in this model in spite of the lower overall muscle volume. Strength training also has potential benefits for injury prevention [53] and could give a competitive edge when a race comes down to a sprint at the end

### Limitations

The results of our simulations are not guaranteed to be applicable to each individual because the specific changes to create an optimized model depend on the body segment dimensions and muscle strengths of the generic model. The body segment geometries, inertial properties, and muscle geometry of the generic model are the synthesis of a series of carefully executed studies [54–57] that resulted in a model to represent an average male of 1.81m and 75.1kg. The muscle volumes of individual muscles in the model are based on detailed measurements of muscles in cadavers of both young [58] and older adults [59]. The specific tension of individual muscles, which scale the muscle volumes to maximal isometric forces, were scaled to match maximal torque-angle relationships established by dynamometer measurements and enable the model to reach realistic vertical jumping heights [56,60]. Since the development of this model, many studies have proven its usefulness for simulations of different tasks. We are aware of specific adaptations to the model to improve simulation of exceptional performance (e.g., elite sprinting [35]) and extreme motions (e.g. deep squats [61]). However, we chose to not implement these adaptations as tuning the model might compromise the behavior in other tasks. Thus, while the generic model has been carefully developed and tested, the specific numerical results presented here depend on its properties. It is possible, however, to overcome this limitation and to understand how performance depends on musculoskeletal parameters for a specific person if a personalized musculoskeletal model is available. Developing personalized models is a challenge for future research.

Another limitation of our study, and an interesting avenue for future work, lies in how we scaled muscle volumes with scaling of the skeleton segments. A larger person will have higher muscle volume than a smaller person with the same fat percentage. However, the assumption used in this study of a linear increase in muscles volume distributed uniformly across muscles influences simulated results. For example, in our simulator, the strength of all muscles increased when the optimizer increased the size of the torso or upper arm. An alternate approach to explore in future work is to assign each muscle to one or more segments and scale muscle volume with the volume changes of the assigned segments.

Our simulations of sprinting have lower hip and knee flexion angles during swing compared to experimental measurements (Supplemental Figure S.3), as observed in prior sprinting simulations [62]. This might be due to passive hip and knee extension moments becoming large in the model at the more extreme flexion angles. This is a known limitation of the model we used, and attempts to mitigate this issue have been performed [61]. However, sprinting simulations of the generic model with passive forces disabled did not result in larger knee flexion angles, thus we decided not to decrease the passive muscle forces since this may compromise the simulation of other motions for which the original model was developed.

The speeds at which we simulated marathon running and that we found for sprinting are below elite level. The lower sprint speed is mainly due to the relative weakness of the muscles in our generic model. When multiplying the maximal isometric force of all muscles in the generic model by a factor of 2, as was done in several previous sprinting simulation studies [32,35], the maximal running speed increased to 11.1m/s, which is closer to the fastest ever recorded speed of 12.42m/s. The speed of 3.33m/s was chosen as a representative speed for the marathon because it has a similar ratio to elite marathon running speed as the maximal running speed of the generic model to elite sprinting speed.

Our optimization is limited to maximizing speed and minimizing energy consumption and ignores other important factors. For example, the segment scaling could potentially lead to very narrow and long bone geometries that might be more prone to injury. Also, the optimal running techniques simulated might induce high force peaks that generate painful joint loads, and we did not limit these forces in the simulations. Although the vertical impulse is similar in simulation and experiment, the simulated vertical ground reaction forces have a higher peak compared to experimental data (Figure S.4).

## Conclusions and Future Directions

Our open-source simulator was able to simultaneously optimize many musculoskeletal parameters in a three-dimensional simulation of running to uncover determinants of sprint and marathon running performance. The simulator could also be used to optimize performance of other athletic tasks such as jumping and accelerative running, which are important for many sporting events. While the present study focused on athletic performance, our approach could also be used in other applications. For example, the simulator could determine minimal strength requirements to safely perform activities of daily living, could guide strength training interventions in elderly people, or estimate the effects of musculoskeletal surgery.

## METHODS

We performed ten predictive simulations of a running gait (overview in Table S.1) using the three-dimensional musculoskeletal model developed by Hamner and colleagues [32], which includes 92 muscle actuators, 8 torque actuators, and 31 degrees of freedom. The first two simulations optimized the excitations of the muscles and torque actuators (i.e., motor coordination) to maximize sprinting and marathon performance. The third and fourth simulations optimized for both motor coordination and segment geometry, which included three scaling factors (one for each dimension) for each body segment in the model. Scaling the segments affects the bone geometry, muscle insertion points, muscle moment arms, and muscle properties, such as optimal fiber length and tendon slack length. As such, these simulations yielded a model with sprint-optimized skeleton body segment dimensions and a model with marathon-optimized body segment dimensions. Importantly, when scaling the skeleton, we chose to scale muscle volumes proportionally to the change in whole body mass and adjust body mass when scaling the body segments by assuming constant density. As such, a heavier model has stronger muscles. Next, we evaluated sprinting and marathon performance for the models with body segment dimensions optimized for the opposite task. Finally, we simulated optimal strengthening, yielding a sprint-optimized muscle strength model and a marathon-optimized muscle strength model. We also evaluated these strength-adjusted models for the opposite task,

### Differentiable musculoskeletal simulator

We developed a differentiable musculoskeletal simulator (Figure 4). Every function within this simulator is differentiable; thus, we can rapidly obtain the gradient of the output with respect to all its inputs using automatic differentiation rather than finite differencing, and all gradients are continuous.

**Figure 4.**
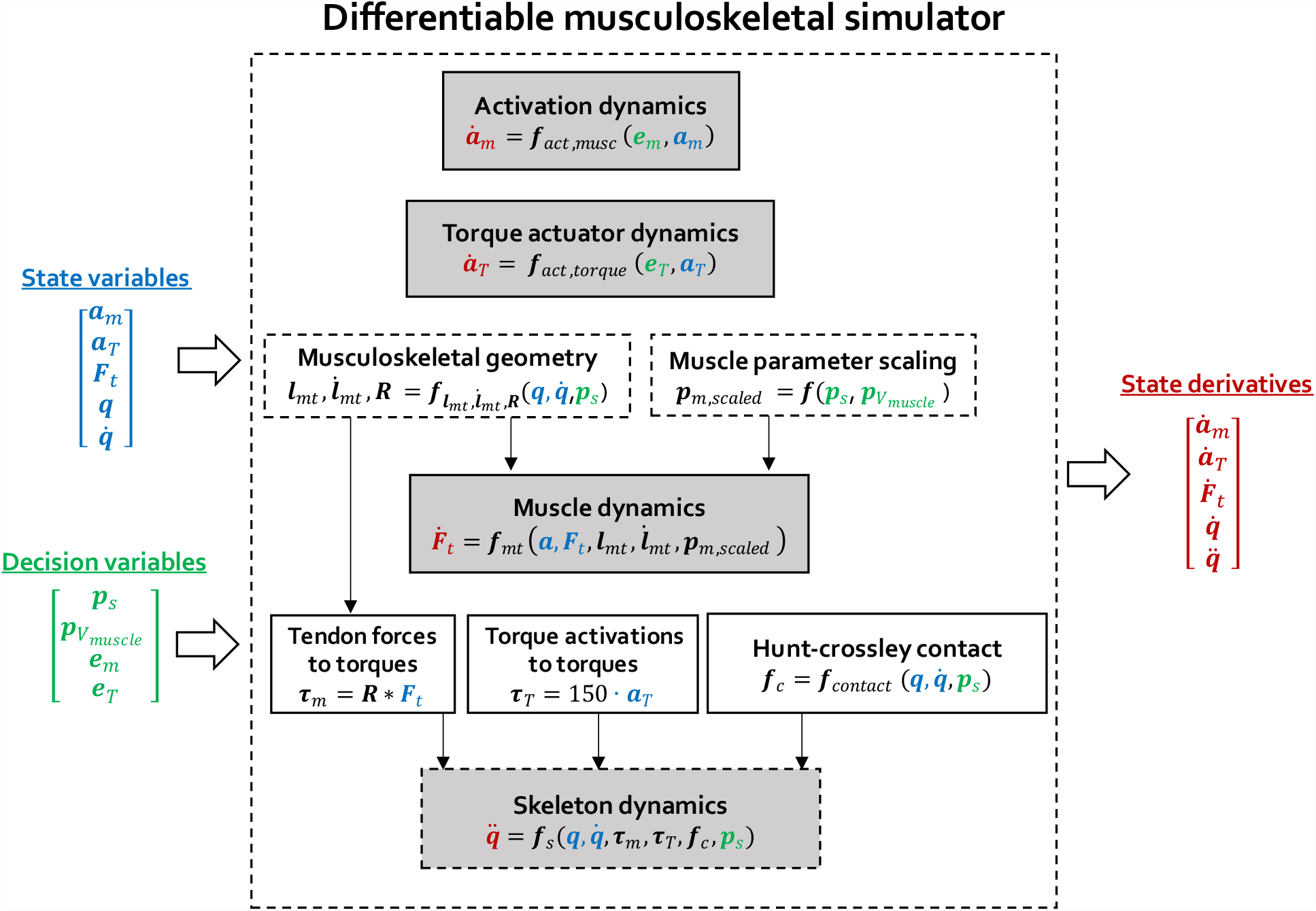
Our differentiable musculoskeletal simulator generates the derivatives of the state variables given the state variables (muscle activations **a**_m_, torque actuator activations **a**_T_, tendon forces **F**_t_, generalized positions **q** and velocities 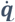) and the decision variables (skeleton segment scaling factors **p**_s_, muscle volume scaling factors 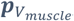, muscle excitations **e**_m_, torque actuator excitations **e**_T_). This is achieved by evaluating a set of dynamics equations: activation dynamics, torque actuator dynamics, muscle dynamics, and skeleton dynamics. Evaluating muscle and skeleton dynamics depends on the outputs of musculoskeletal geometry computations (i.e., muscle-tendon lengths **l**_mt_ and velocities 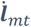 and muscle moment-arm matrices **R**) and on the scaled muscle parameters (**p**_m,scaled_). Since the scaling of the skeleton and muscle volumes are decision variables, we formulated musculoskeletal geometry computation, muscle parameter scaling and skeleton dynamics as a differentiable function of these decision variables. The dotted boxes were the parts of the simulator where we turned the state-of-the-art from non-differentiable to differentiable computation. Tendon forces are mapped to joint muscle torques (**τ**_m_) by the moment-arm matrix (**R**). Torque actuator activations are scaled to torque actuator torques (**τ**_T_) by a scaling factor of 150 [63]. A contact function (**f**_contact_) based on the Hunt-Crossley contact model gives the generalized forces resulting from contact (**f**_c_).

At each timestep of a simulation our musculoskeletal simulator gives the state derivatives given the state and decision variables. The state (***x*** *∈* ℝ^**254**^) of the musculoskeletal simulator is determined by the activations (***a***_*m*_ *∈* ℝ^**92**^) of the 92 included lower limb muscles, the force in each tendon (***F***_*t*_ *∈* ℝ^**92**^), the activation of the eight torque actuators of the upper limb degrees of freedom (***a***_*T*_ *∈* ℝ^**8**^), and the generalized positions (***q*** *∈* ℝ^**31**^) and velocities 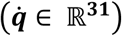, including the six degrees-of-freedom of the pelvis and 25 joint angles:

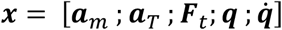

The decision variables that form the input to the musculoskeletal simulator at each timestep are the muscle (***e***_*m*_ *∈* ℝ^**92**^) and torque actuator excitations (***e***_*T*_ *∈* ℝ^**8**^), the scaling factors of the skeleton segments ***p***_*s*_ ∈ ℝ^3×18^) and scaling factors of the muscle volumes 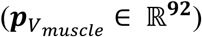.

The state derivatives are described by muscle activation dynamics [64]:

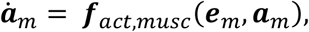

muscle-tendon dynamics [64]:

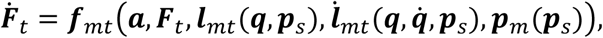

torque actuator activation dynamics [8]:

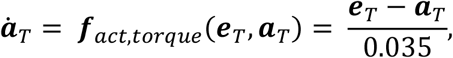

and skeleton dynamics [40]:

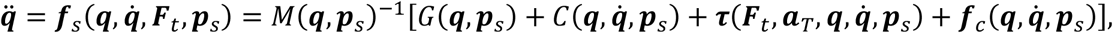

where ***e***_*m*_ are the muscle excitations, ***l***_*mt*_ muscle-tendon lengths, 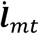 muscle-tendon velocities, ***p****s* the skeleton parameters, ***p***_*m*_ the muscle parameters, ***e****T* the torque actuator excitations, ***τ*** biological joint torques, ***f***_*c*_ the function describing the generalized forces that result from contact, *M* the mass matrix, *G* the vector of gravitational forces, and *C* the vector of Coriolis and centrifugal forces. Contact is modelled using a Hunt-Crossley model and occurred between eight contact spheres attached to each of the feet and the ground. The location and properties of these contact spheres are as described in [36].

The biological joint torques are the result of the torques generated by the muscles ***τ***_*m*_), the torques generated by the torque actuators for the upper limbs ***τ***_*T*_) and passive joint torques ***τ***_*pas*_):

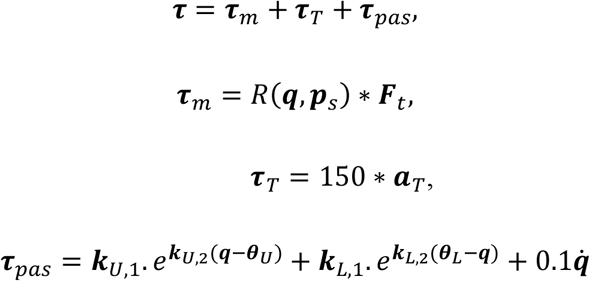

The muscle torques result from matrix multiplication between the muscle forces with ***R***(***q, p***_*s*_) the 92×31 matrix of moment arms of the muscles with respect to the joints. The torque actuator torques result from scaling the torque actuator activations that are bounded between -1 and 1 with 150 as in [35,63].

The passive joint torques (***τ***_*pas*_) consist of joint limit torques to model ligaments for the muscle driven joints and damper joint torques for all joints with a damping constant of 0.1 Nm.s/rad. The joint limit torques are parametrized by six parameters (***k***_*U*,1_, ***k***_*U*,2_, ***θ***_*U*_, ***k***_*L*,1_, ***k***_*L*,2_, ***θ***_*L*_) describing the exponential decay and increase across the range of motion for every joint coordinate. The parameters are taken from [56].

Finally, for the metatarsophalangeal joint, which is not muscle actuated, a spring-damper joint torque is added that has different parameters depending on whether sprinting (stiffness: 40 Nm/rad, damping: 0.4 Nm.s/rad) [35] or marathon running (stiffness: 25 Nm/rad, damping: 1.9 Nm.s/rad) [36] is simulated.

The skeleton parameters ***p***_*s*_ ∈ ℝ^3*x*18^ consist of three scaling factors for each body segment, one for scaling each dimension. When scaling the model, we assume the following 18 bodies: the pelvis, trunk+head, plus two of each of the talus, calcaneus, toes, shank, thigh, upper arms, lower arms, hands.

To simplify the problem, we use constraints to impose symmetry and to require identical scaling factors for the trunk+head, upper arms, lower arms, and hands, and identical scaling factors for talus, calcaneus and toes. Importantly, scaling the body segments affects the following quantities in the skeleton dynamics: *M, G, C*, ***f***_*c*_, defined above.

The skeleton segment scaling factors are also an input to the musculoskeletal geometry computation that calculates the muscle-tendon lengths, muscle-tendon velocities and moment-arm matrix as a function of the generalized coordinates, generalized velocities and skeleton segment scaling factors:

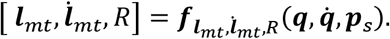

The Hill-type muscle parameters ***p***_*m*_ consist of: the muscle physiological cross sectional area (***PCSA***), the specific tension of muscle fibers (***σ***), tendon slack length (***l***_*T,s*_), optimal fiber length (***l***_*m,opt*_), pennation angle (***α***_*m*_) and tendon stiffness (***k***_*T*_) [64]. The cross-sectional area and specific tension determines the muscle maximal isometric force:

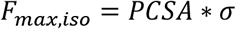

From the PCSA and the optimal fiber length, the muscle volume is calculated:

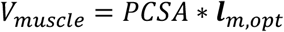

which is an input to several other computations including the metabolic energy consumption [65].

To mimic strength training, we allowed *V*_*muscle*_ of individual muscles to be scaled by a scaling factor 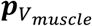:

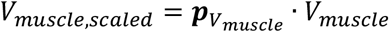

and as such changing the muscle maximal isometric force.

When scaling the skeleton, the tendon slack length and optimal fiber length are adapted as well depending on the total length change of the muscle tendon unit length when the model is placed in the anatomical pose (***q*** = **0**):

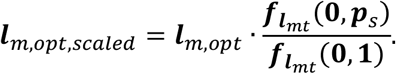

The same ratio is applied to scale tendon slack length.

Scaled muscle parameters are thus a function of original muscle parameters, the scaling factors of the skeleton segments and the scaling factor for the muscle volume:

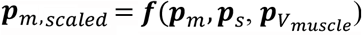

#### Turning non-differentiable into differentiable computation

Optimizing the skeleton body segment dimensions is not feasible using the state-of-the-art musculoskeletal simulator OpenSim [37] and Simbody [40] due to its non-differentiable computation. The simulator from Falisse et al. [8] enables differentiable computation for a part of the OpenSim-Simbody simulator and serves as a starting point for our simulator. However, the simulator of from Falisse et al. [8]is not differentiable with respect to all musculoskeletal variables of interest. As such, an important technical contribution of this work is to make the entire musculoskeletal simulator differentiable. We identified two non-differentiable operations when relying on OpenSim and Simbody. First, musculoskeletal geometry computations, 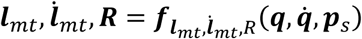, are non-differentiable with respect to joint coordinates (***q***), velocities 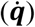 and skeleton segment scaling factors (***p***_*s*_), and the first-order derivatives are not guaranteed to be continuous. Second, the skeleton dynamics, 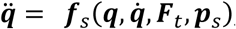, are non-differentiable with respect to ***p***_*s*_, which determines the contributions of *M, G, C*, ***f***_*c*_ to the skeleton dynamics.

### Differentiable musculoskeletal geometry computation

We implemented the musculoskeletal geometry computation as a differentiable neural network function:

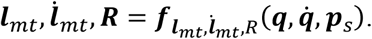

Musculoskeletal geometry computation in OpenSim is executed as follows: first, based on the skeleton segment scaling factors the bone geometries, muscle insertion points, muscle via points and muscle wrapping surfaces are adapted,) next, using the scaled geometry the muscle-tendon lengths and moment arms are calculated. Both parts implemented in OpenSim as non-differentiable operations with non-continuous first-order derivatives.

To resolve this, we implemented a shallow (two hidden layers) neural network to calculate ***l***_*mt*_, ***R*** for every muscle. Having a differentiable function for the computation of ***l***_*mt*_ for each pose (***q***) and set of skeleton segment scaling factors (***p***_*s*_) allows us to have a differentiable function to perform muscle parameter scaling 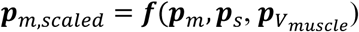. Finally, the muscle-tendon velocities are computed using the chain rule:

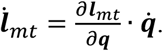

Having a separate neural network for every muscle reduces computational complexity, as we can take into account that every muscle only attaches to a limited number of bones.

We used OpenSim to generate training data and started from the generic OpenSim model for running[32]. We scaled 2,000 versions of this model using the OpenSim Scale tool [37]. The scaling factors, that serve as input to the Scale Tool, are a 54×1 vector (representing the three dimensions of the 18 bodies of the model), of which each element is drawn from a uniform distribution: (*U*.8,1.2). We chose the respective lower and upper limits of 0.8 and 1.2 as these cover most of the variation in an adult population.

With the scaled models at our disposal, we generated training samples for all the different muscles. For every training sample, we randomly selected one of the 2000 models and randomly drew a joint pose from a uniform distribution that has lower and upper limits according to the joint range of motion. We put the model in that joint pose and observed the muscle tendon length and moment arm with respect to the joint coordinates that are actuated by the muscle. Depending on whether the muscle actuates, one, two or more joint coordinates, we drew 20,000, 82,000, or 200,000 samples for that muscle.

We then trained a separate neural network for each muscle using Adam [66] with a mean-squared-error loss over 1,000 epochs and a batch size of 64. We used feedforward neural networks with ‘tanh’ activation functions and two hidden layers. The size of the hidden layers was 8, 12, or 16 depending on whether the muscle actuates 1, 2 or more generalized coordinates.

We experimented to minimize the size of the neural networks to reduce computational complexity without sacrificing accuracy of the approximation. With the described set-up we confirmed that our predictive simulations of walking and running at 3.33m/s yielded kinematics and muscle activation very close to the those with the original OpenSim model and for OpenSim models with all scaling factors at 0.85 or 1.15.

### Differentiable skeleton dynamics

Skeleton dynamics are based on SimBody [40]. We adapted the source code transformation tool from [67] to enable automatic differentiation of the skeleton dynamics with respect to the segment scaling parameters ***p***_*s*_. This source code transformation tool analyzes a given function’s source code and outputs the gradient of that function. The source code transformation tool takes a customized.cpp description of the SimBody skeleton model and its skeleton dynamics as a function to analyze and differentiate. The tool also requires a description of the variables with respect to which it will generate the gradient. In addition to the state-of-the-art implementation where the functions generalized coordinates, velocities, and accelerations as differentiable input, we extended its functionality to be differentiable with respect to the geometrical scaling factors of each segment. Therefore, we added functions to define how these geometrical segment scaling factors change the mass, center-of-mass location, and inertial properties for every segment. Next, we added functions to define how these segment scaling factors change rotational and translational offsets across the kinematic tree of the skeleton. These functions mimicked how segment scaling factors affect these model properties when using the OpenSim Scale tool, but in a smooth and differentiable way.

Each predictive simulation was solved as a trajectory optimization problem. We simulated steady-state gait (running and sprinting) and assumed symmetry. As such we only needed to simulate half a gait cycle while imposing symmetry, as well as continuity and periodicity constraints for the appropriate states.

#### Trajectory optimization

For every trajectory optimization problem, we optimized at least the motor coordination, consisting of muscle (***e***_*m*_) and torque excitations (***e***_*T*_), and the initial state of the musculoskeletal system (*x (t*_0_)).

For simulations where we optimized either body segment dimensions or muscle volume scaling to maximize task performance, we solved a trajectory optimization problem where either the skeleton scaling parameters (***p***_*s*_) or the scaling of muscle volumes 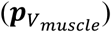 were added to the optimization variables.

### Objective function

The minimal energy objective for marathon running was adapted from Falisse et al. [8] and consists of five main contributions:

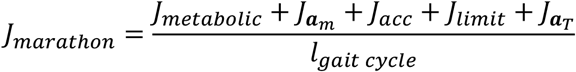

with *J*_*metabolic*_ the muscle metabolic energy based on the Bhargava model [65] of metabolic energy expenditure, 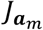 the sum of squared muscle activations modelling muscle fatigue, *J*_*acc*_ the sum of squared joint accelerations modelling motion smoothness, *J*_*limit*_ the sum of squared limit joint torques modelling avoidance of ligament strain, and 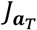 the sum of squared upper limb torque actuator activations modelling upper body fatigue and energy expenditure. The sum of these terms is normalized by the length of the gait cycle. The metabolic cost of running a marathon, which is reported in the results as marathon performance is computed by multiplying 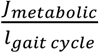 with the length of a marathon: 42,196 m.

A straightforward sprint objective to maximize the average velocity was chosen:

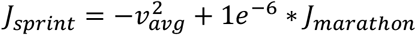

with a small contribution of the marathon energy term to improve the numerical condition of the optimization problem.

### Constraints and bounds

Muscle excitations and activations are bound to be between 0 and 1, whereas torque actuator excitations are bounded between -1 and 1. We include path constraints to avoid penetration between body segments and additional bounds for the joint ranges of motion. These ranges of motion are generous and typically not reached for most degrees of freedom as the modelled passive forces representing ligaments provide a physical joint limit. However, for the upper limb degrees of freedom we use a more strict and representative range of motion that was reached in some simulations. This choice was made as we did not model muscular or ligamentous structures to limit these. Similarly, hip inward (−10°) and outward rotation (+10°) as well as knee extension (0°) bounds were typically reached. For the remaining variables (muscle lengths, joint velocities, muscle velocities, joint accelerations) we use generous bounds that were there to improve numerical stability during optimization and were not reached.

For the simulations of marathon running we imposed the average speed to be 3.33m/s.

For simulations where the skeleton scaling parameters, ***p***_*s*_, were optimized, these were bounded between 0.8 and 1.2. We also imposed a constraint on the Body Mass Index to be between 17.5 and 25.5 to represent a healthy person.

When simulating strength training, the increase of individual muscle physiological cross-sectional areas was limited to 20% and the total increase in muscle volume summed over all muscles was limited to 5% of the initial total muscle volume.

### Direct collocation and implicit dynamics

To improve numerical conditioning, we formulated muscle and skeleton dynamics with implicit rather than explicit differential equations. We therefore introduced derivatives of tendon force and coordinate accelerations as additional controls, and we imposed the nonlinear dynamic equations describing muscle contraction and skeleton dynamics as algebraic constraints.

We used direct collocation to transcribe each trajectory optimization problem into a large sparse nonlinear program. We used a third-order Radau quadrature collocation scheme with 50 mesh intervals per half gait cycle and solved the resulting NLP with the solver IPOPT. All gradients were computed using automatic differentiation, where we relied on CasADi [68].

Because we fix the number of mesh intervals for every simulation problem, we made the mesh interval length a variable to accommodate for different possible stride lengths at a given speed.

## SUPPLEMENTARY MATERIALS

### Overview of the predictive musculoskeletal simulations

We performed ten simulations or sprinting and marathon running (Table S.1). For all simulations the motor coordination was optimized for the task of interest. In some cases the body segment dimensions or the muscle volume were optimized (i.e., were decision variables), in which case, new models were generated.

**Table S.1.** Summary of the ten predictive simulations of marathon running and sprinting.

|  | **Task** | **Motor coordination** | **Body segment scaling factor** | **Muscle volume scaling factor** | **Yields new model?** |
| --- | --- | --- | --- | --- | --- |
| 1 | Sprinting | decision variable | generic | decision variable | - |
| 2 | Marathon | decision variable | generic | decision variable | - |
| 3 | Sprinting | decision variable | decision variable | decision variable | sprint-optimized body segments |
| 4 | Marathon | decision variable | decision variable | decision variable | marathon-optimized body segments |
| 5 | Marathon | decision variable | sprint-optimized body segments | decision variable | - |
| 6 | Sprinting | decision variable | marathon-optimized body segments | decision variable | - |
| 7 | Sprinting | decision variable | decision variable | free | sprint-optimized muscle volume |
| 8 | Marathon | decision variable | decision variable | free | marathon-optimized muscle volume |
| 9 | Marathon | decision variable | decision variable | sprint-optimized muscle strength | - |
| 10 | Sprinting | decision variable | decision variable | marathon-optimized muscle strength | - |

### Comparing predicted kinematics to experimental kinematics

Our simulated kinematics represent many of the key features of experimentally measured kinematics during running and sprinting. For marathon running, which we defined as running at 3.33m/s, we compared simulated kinematics and ground reaction forces to experimental data from ten individuals running at about 3.1m/s [32] (Figure S.1, S.2). The ankle, hip flexion, and hip adduction kinematics are similar. Our simulated hip rotation kinematics fall within the variability observed in experimental hip rotation kinematics. The knee kinematics have a similar waveform, but simulations show lower peak knee flexion during swing and higher knee flexion during stance. The pelvis and lumbar degrees of freedom have similar waveforms, but in the simulations there is a more limited range of motion. The pelvis posterior tilt is lower in simulation compared to the experimental data. This explains a similar but opposite offset in the lumbar extension angle and hip flexion angle, meaning that segment orientations for the torso and femurs do not have an offset between experiments and simulation.

**Figure S.1.**
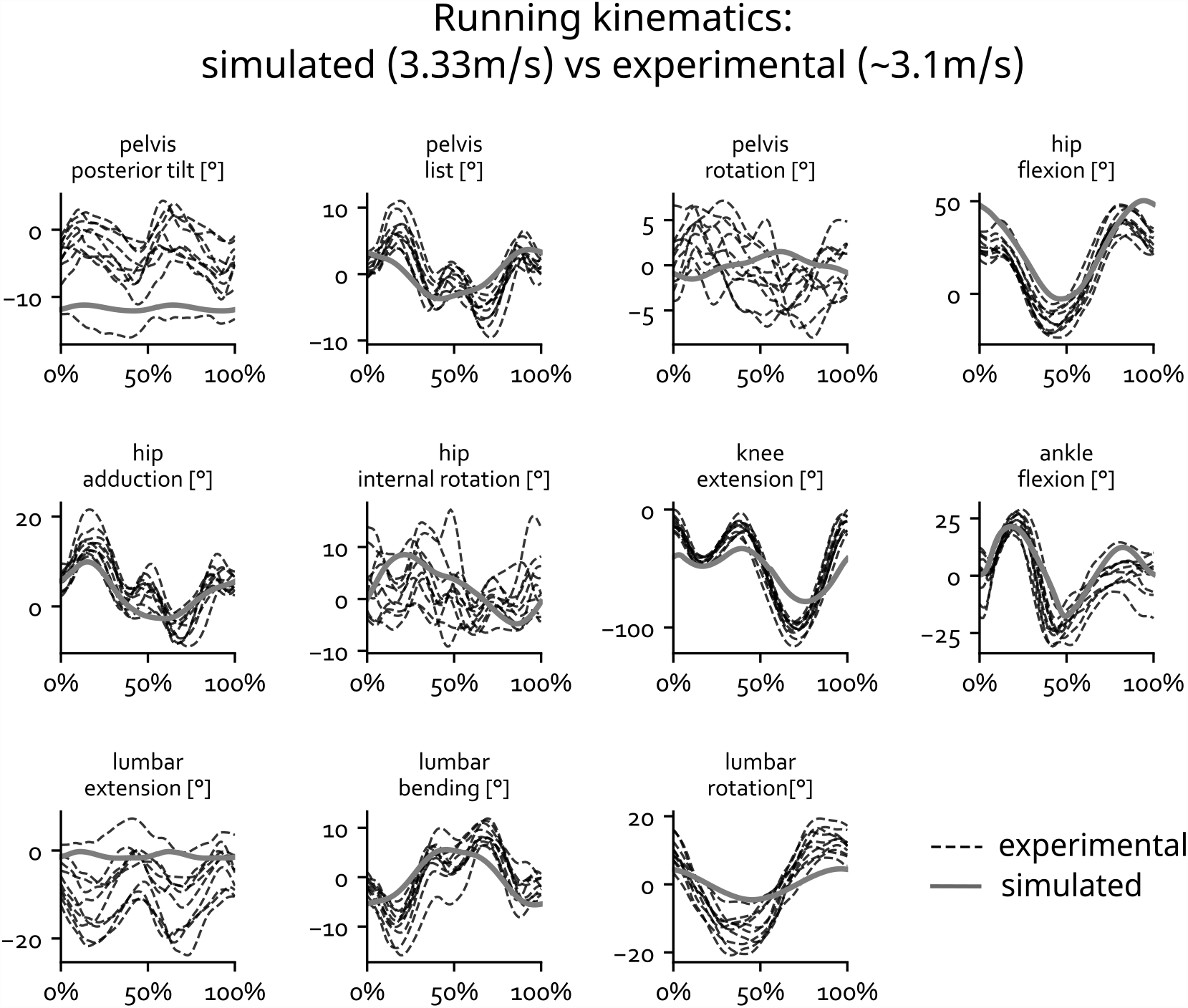
Simulated (solid line) and experimental (dotted lines) kinematics during running. The gait cycle is defined from foot strike to foot strike.

Simulated and experimental ground reaction forces were also similar (Figure S.2). In our simulations the stance phase was slightly longer than in the experiments.

**Figure S.2.**
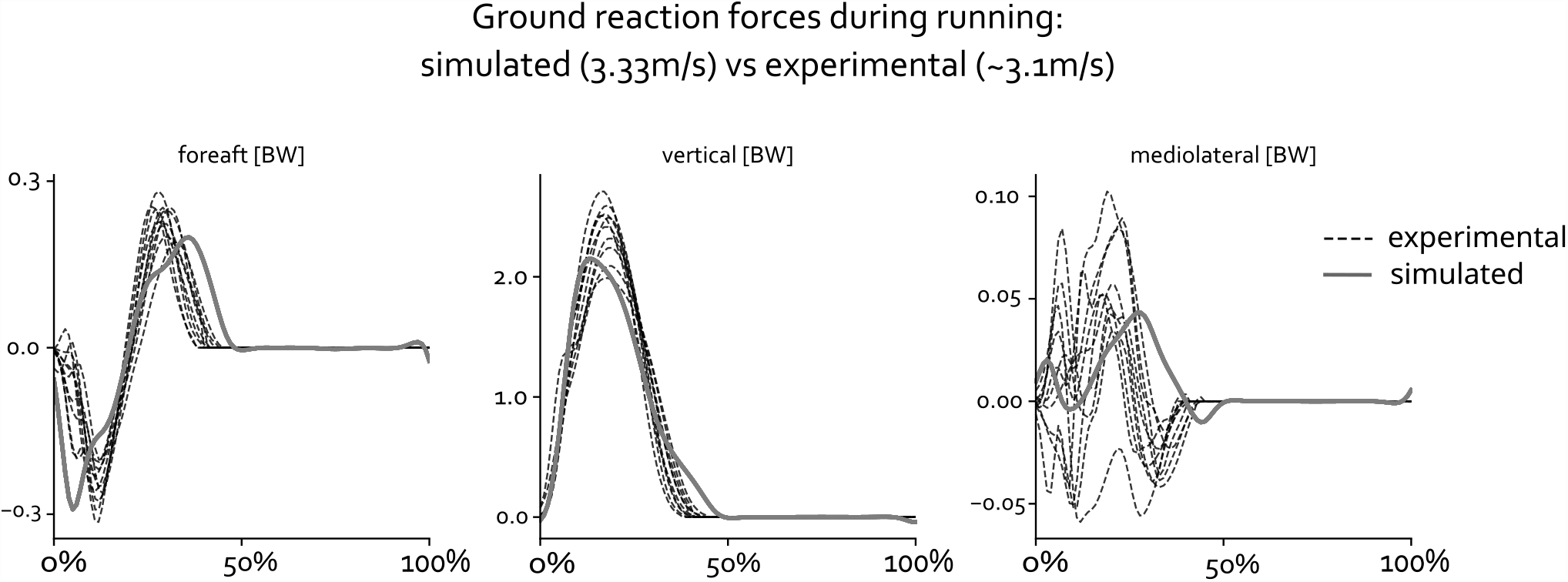
Simulated (solid line) and experimental (dotted lines) ground reaction forces during running. The gait cycle is defined from foot strike to foot strike.

We compared our sprinting simulation to a sprinting gait cycle from an individual running at about 9.45m/s [34] (Figure S.3). The most important differences between simulated and experimental kinematics are a lower simulated peak knee and hip flexion during swing. This has been observed in other simulations studies as well [62]. These differences may be due to the fact that (1) we did not tune any of the muscle parameters (e.g., to reduce passive forces), which might prevent such high values for knee and hip flexion [61], and (2) our model has a lower sprint speed than in the experiment.

The shape of the ground reaction force and duration of contact are similar in simulation and the experiment (Figure S.4). The simulated vertical ground reaction force reaches a significantly higher peak than observed in the experiment. The contact impulse is similar for experiment and simulation (not shown explicitly) as the simulated higher peak is shorter than the experimental one. Such simulated running technique might be painful in reality because of high impact forces and therefore not seen in the experiment. Numerical parameters of contact models to represent real foot-ground contact are hard to determine. The parameters we used were calibrated against experimental data in a prior study [36], but due to the highly non-linear characteristics of contact physics large changes can occur when the model behaves outside of the experimental data for which it was calibrated. The fore-aft ground reaction force is smaller in simulation for both the braking and acceleration component.

**Figure S.3.**
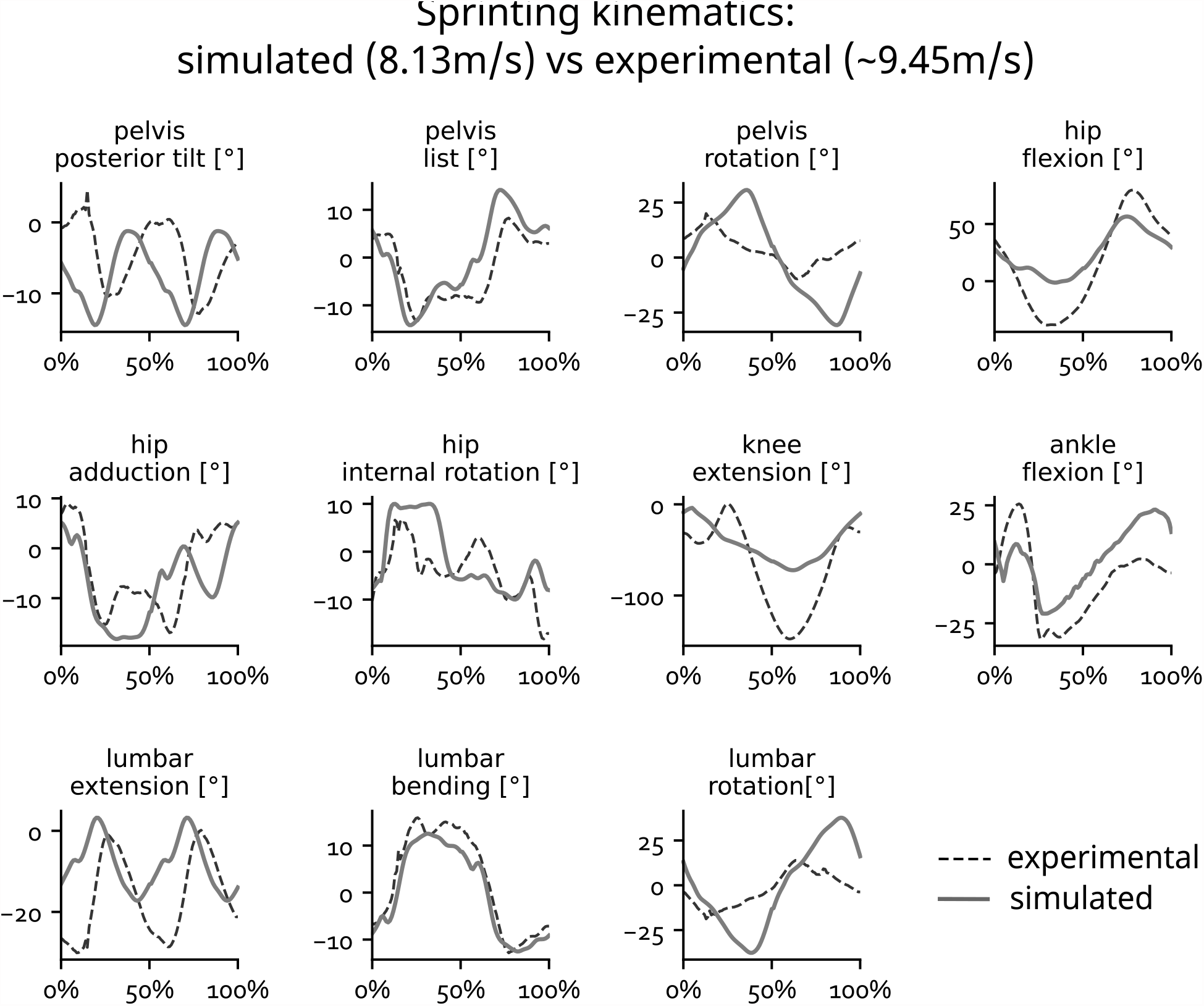
Simulated (solid line) and experimental (dotted lines) kinematics during sprinting. The gait cycle is defined from foot strike to foot strike.

**Figure S.4.**
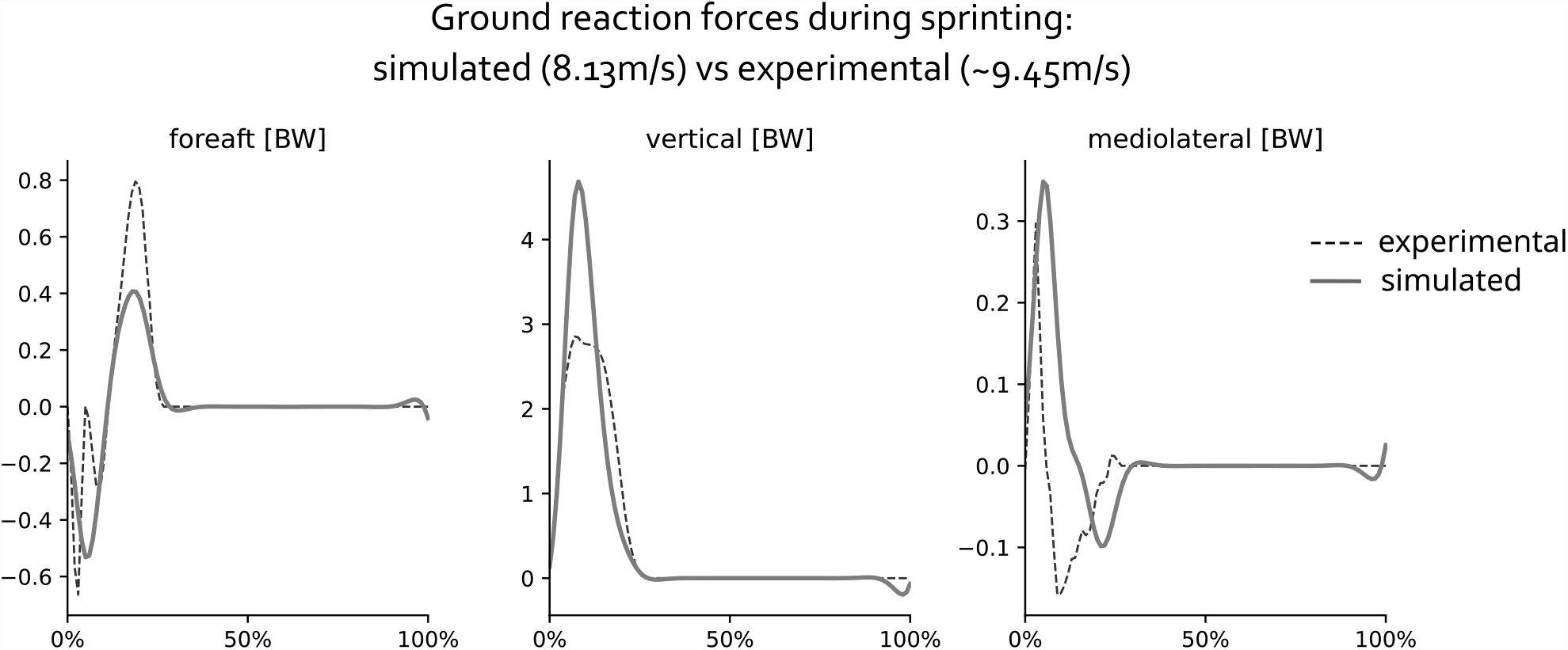
Simulated (solid line) and experimental (dotted lines) ground reaction forces during sprinting. The gait cycle is defined from foot strike to foot strike.

### Validating muscle-tendon length moment arm and approximation

Our neural network approximates the muscle-tendon lengths and moment arms within 2mm for all poses during a sprinting cycle and across all optimized models (Figure S.5). For each model, we calculated for each muscle the average absolute error between ground truth and our approximation of muscle-tendon length and moment arm during sprinting. The mean errors are well below 1mm.

**Figure S.5.**
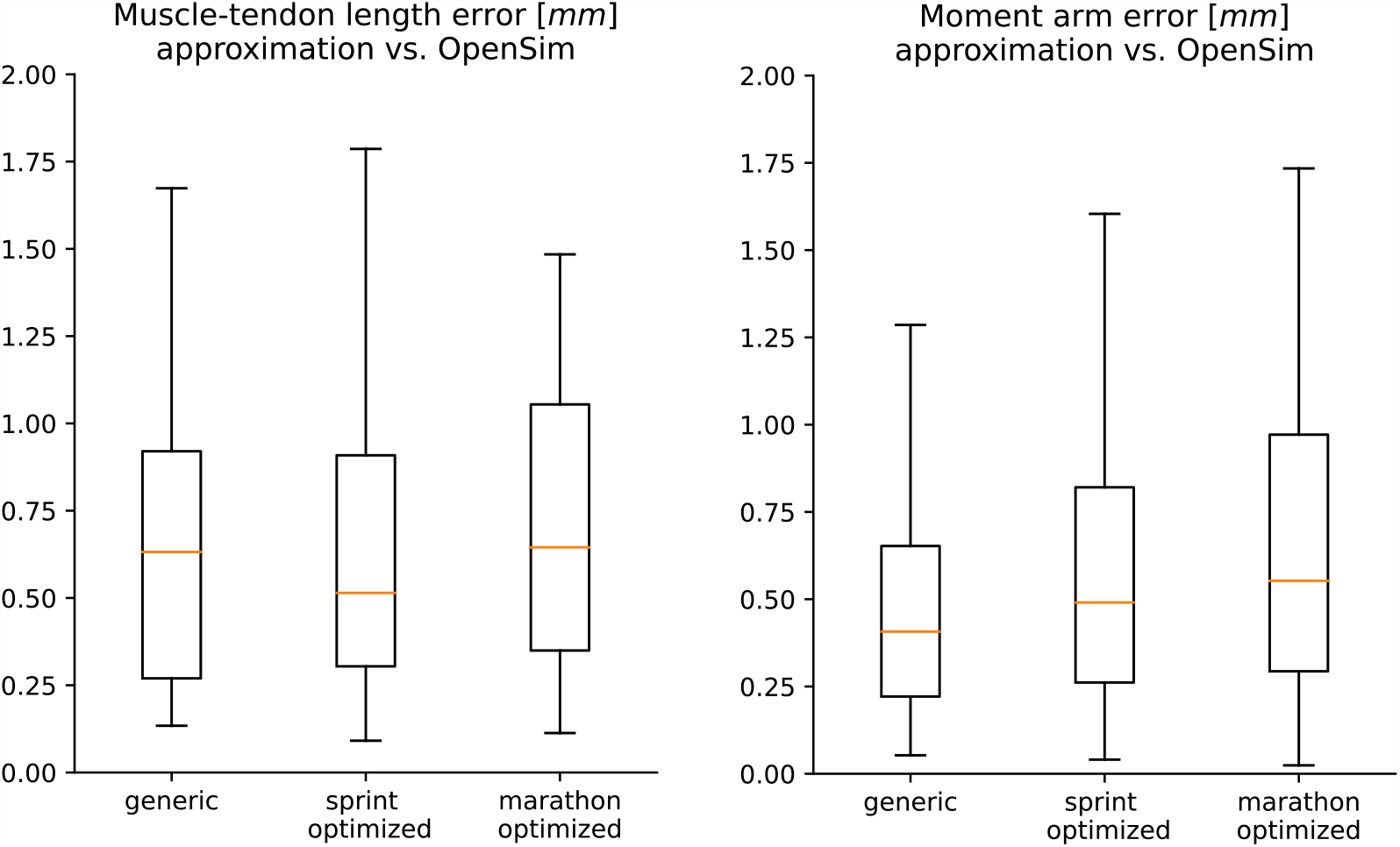
Muscle-tendon length and moment arm error. We show average absolute errors across all the muscles.

## REFERENCES

1. Sedeaud A, Marc A, Marck A, Dor F, Schipman J, Dorsey M, et al. BMI, a Performance Parameter for Speed Improvement. PLoS One. 2014;9: e90183. doi:10.1371/JOURNAL.PONE.0090183

2. Hawes MR, Sovak D. Morphological prototypes, assessment and change in elite athletes. J Sports Sci. 1994;12: 235–242. doi:10.1080/02640419408732168

3. Foley JP, Bird SR, White JA. Anthropometric comparison of cyclists from different events. Br J Sport Med. 1989;23: 30–3. doi:10.1136/bjsm.23.1.30

4. Bobbert MF, Van Zandwijk JP. Sensitivity of vertical jumping performance to changes in muscle stimulation onset times: a simulation study. Biol Cybern 1999 812. 1999;81: 101–108. doi:10.1007/S004220050547

5. Wilson C, Yeadon MR, King MA. Considerations that affect optimised simulation in a running jump for height. J Biomech. 2007;40: 3155–3161. doi:10.1016/j.jbiomech.2007.03.030

6. Miller RH, Umberger BR, Caldwell GE. Limitations to maximum sprinting speed imposed by muscle mechanical properties. J Biomech. 2012;45: 1092–1097. doi:10.1016/J.JBIOMECH.2011.04.040

7. Dembia CL, Bianco NA, Falisse A, Hicks JL, Delp SL. OpenSim Moco: Musculoskeletal optimal control. doi:10.1101/839381

8. Falisse A, Serrancolí G, Dembia CL, Gillis J, Jonkers I, De Groote F. Rapid predictive simulations with complex musculoskeletal models suggest that diverse healthy and pathological human gaits can emerge from similar control strategies. J R Soc Interface. 2019;16. doi:10.1098/RSIF.2019.0402

9. Di Salvo V, Baron R, Tschan H, Calderon Montero FJ, Bachl N, Pigozzi F. Performance characteristics according to playing position in elite soccer. Int J Sports Med. 2007;28: 222–227. doi:10.1055/S-2006-924294/ID/35

10. Faude O, Koch T, Meyer T. Straight sprinting is the most frequent action in goal situations in professional football. https://doi.org/101080/026404142012665940. 2012;30: 625–631. doi:10.1080/02640414.2012.665940

11. Hoogkamer W, Kram R, Arellano CJ. How Biomechanical Improvements in Running Economy Could Break the 2-hour Marathon Barrier. Sport Med. 2017;47: 1739–1750. doi:10.1007/S40279-017-0708-0/FIGURES/9

12. Joyner MJ, Ruiz JR, Lucia A. The two-hour marathon: who and when? J Appl Physiol. 2011;110: 275–277. doi:10.1152/japplphysiol.00563.2010

13. Uth N. Anthropometric Comparison of World-Class Sprinters and Normal Populations. J Sports Sci Med. 2005;4: 608. Available: /pmc/articles/PMC3899678/

14. Barbieri D, Zaccagni L, Babić V, Rakovac M, Mišigoj-Duraković M, Gualdi-Russo E. Body composition and size in sprint athletes. J Sports Med Phys Fitness. 2017;57: 1142–1146. doi:10.23736/S0022-4707.17.06925-0

15. Tomita D, Suga T, Terada M, Tanaka T, Miyake Y, Ueno H, et al. A pilot study on a potential relationship between leg bone length and sprint performance in sprinters; are there any eventrelated differences in 100-m and 400-m sprints? BMC Res Notes. 2020;13. doi:10.1186/S13104-020-05140-Z

16. Miller R, Balshaw TG, Massey GJ, Maeo S, Lanza MB, Johnston M, et al. The Muscle Morphology of Elite Sprint Running. Med Sci Sports Exerc. 2021;53: 804–815. doi:10.1249/MSS.0000000000002522

17. Sharma SS, Dixit NK. Somatotype of athletes and their performance. Int J Sports Med. 1985;6: 161–162. doi:10.1055/S-2008-1025831/BIB

18. Norton K, Olds T. Morphological evolution of athletes over the 20th century: Causes and consequences. Sport Med. 2001;31: 763–783. doi:10.2165/00007256-200131110-00001/FIGURES/15

19. Carter J. The somatotypes of athletes--a review. Hum Biol. 1970;42: 535–569.

20. Blazevich AJ. Predicting Sprint Running Times From Isokinetic and Squat Lift Tests: A Regression Analysis. J Strength Cond Res. 1998 [cited 21 Oct 2022]. Available: https://www.academia.edu/29201982/Predicting_Sprint_Running_Times_From_Isokinetic_and_Squat_Lift_Tests_A_Regression_Analysis

21. Muñoz CS, Muros JJ, Belmonte ÓL, Zabala M. Anthropometric Characteristics, Body Composition and Somatotype of Elite Male Young Runners. Int J Environ Res Public Heal 2020, Vol 17, Page 674. 2020;17: 674. doi:10.3390/IJERPH17020674

22. Laumets R, Viigipuu K, Mooses K, Mäestu J, Purge P, Pehme A, et al. Lower Leg Length is Associated with Running Economy in High Level Caucasian Distance Runners. J Hum Kinet. 2017;56: 229. doi:10.1515/HUKIN-2017-0040

23. Ueno H, Suga T, Takao K, Miyake Y, Terada M, Nagano A, et al. The potential relationship between leg bone length and running performance in well-trained endurance runners. J Hum Kinet. 2019;70: 165–172. doi:10.2478/HUKIN-2019-0039

24. Mooses M, Mooses K, Haile DW, Durussel J, Kaasik P, Pitsiladis YP. Dissociation between running economy and running performance in elite Kenyan distance runners. J Sports Sci. 2015;33: 136–144. doi:10.1080/02640414.2014.926384

25. Davismes L, Levinrad I. An evaluation of hamstring / quadricep strength ratios in elite long distance runners and sprinters. South African J Sport Med. 1998;5. doi:10.10520/AJA10155163_990

26. Deane RS, Chow JW, Tillman MD, Fournier KA. Effects of hip flexor training on sprint, shuttle run, and vertical jump performance. J strength Cond Res. 2005;19: 615–621. doi:10.1519/14974.1

27. Lee SSM, Piazza SJ. Built for speed: musculoskeletal structure and sprinting ability. J Exp Biol. 2009;212: 3700–3707. doi:10.1242/JEB.031096

28. King MA, Kong PW, Yeadon MR. Maximising forward somersault rotation in springboard diving. J Biomech. 2019;85: 157–163. doi:10.1016/J.JBIOMECH.2019.01.033

29. McErlain-Naylor SA, King MA, Felton PJ. A review of forward-dynamics simulation models for predicting optimal technique in maximal effort sporting movements. Appl Sci. 2021;11: 1–20. doi:10.3390/app11041450

30. Miller RH, Umberger BR, Caldwell GE. Sensitivity of maximum sprinting speed to characteristic parameters of the muscle force–velocity relationship. J Biomech. 2012;45: 1406–1413. doi:10.1016/J.JBIOMECH.2012.02.024

31. Miller RH, Umberger BR, Caldwell GE. Limitations to maximum sprinting speed imposed by muscle mechanical properties. J Biomech. 2012;45: 1092–1097. doi:10.1016/J.JBIOMECH.2011.04.040

32. Hamner SR, Seth A, Delp SL. Muscle contributions to propulsion and support during running. J Biomech. 2010;43: 2709–2716. doi:10.1016/J.JBIOMECH.2010.06.025

33. Growth Charts - Homepage. [cited 28 Jul 2023]. Available: https://www.cdc.gov/growthcharts/

34. Dorn TW, Schache AG, Pandy MG. Muscular strategy shift in human running: dependence of running speed on hip and ankle muscle performance. J Exp Biol. 2012;215: 1944–1956. doi:10.1242/JEB.064527

35. Haralabidis N, Serrancolí G, Colyer S, Bezodis I, Salo A, Cazzola D. Three-dimensional data-tracking simulations of sprinting using a direct collocation optimal control approach. PeerJ. 2021;9: e10975. doi:10.7717/peerj.10975

36. Falisse A, Afschrift M, De Groote F. Modeling toes contributes to realistic stance knee mechanics in three-dimensional predictive simulations of walking. PLoS One. 2022;17: e0256311. doi:10.1371/JOURNAL.PONE.0256311

37. Seth A, Hicks JL, Uchida TK, Habib A, Dembia CL, Dunne JJ, et al. OpenSim: Simulating musculoskeletal dynamics and neuromuscular control to study human and animal movement. Schneidman D, editor. PLOS Comput Biol. 2018;14: e1006223. doi:10.1371/journal.pcbi.1006223

38. Nocedal J, Wright SJ. Numerical optimization. Springer Ser Oper Res Financ Eng. 2006; 1–664. doi:10.1201/b19115-11

39. Groote F De, Demeulenaere B, Swevers J, Schutter J De, Jonkers I, Groote F De, et al. A physiology-based inverse dynamic analysis of human gait using sequential convex programming: a comparative study. Comput Methods Biomech Biomed Engin. 2012;15: 1093–1102. doi:10.1080/10255842.2011.571679

40. Sherman MA, Seth A, Delp SL. Simbody: multibody dynamics for biomedical research. Procedia IUTAM. 2011;2: 241–261. doi:10.1016/J.PIUTAM.2011.04.023

41. Hunt KH, Crossley FRE. Coefficient of Restitution Interpreted as Damping in Vibroimpact. J Appl Mech. 1975;42: 440–445. doi:10.1115/1.3423596

42. Dowson MN, Nevill ME, Lakomy HKA, Nevill AM, Hazeldine RJ. Modelling the relationship between isokinetic muscle strength and sprint running performance. https://doi.org/101080/026404198366786. 2011;16: 257–265. doi:10.1080/026404198366786

43. Newman MA, Tarpenning KM, Marino FE. Relationships between isokinetic knee strength, single-sprint performance, and repeated-sprint ability in football players. J strength Cond Res. 2004;18: 867–872. doi:10.1519/13843.1

44. Handsfield GG, Knaus KR, Fiorentino NM, Meyer CH, Hart JM, Blemker SS. Adding muscle where you need it: non-uniform hypertrophy patterns in elite sprinters. Scand J Med Sci Sports. 2017;27: 1050–1060. doi:10.1111/SMS.12723

45. Ravn P, Cizza G, Bjarnason NH, Thompson D, Daley M, Wasnich RD, et al. Low Body Mass Index Is an Important Risk Factor for Low Bone Mass and Increased Bone Loss in Early Postmenopausal Women. J Bone Miner Res. 1999;14: 1622–1627. doi:10.1359/JBMR.1999.14.9.1622

46. Tenforde AS, Carlson JL, Chang A, Sainani KL, Shultz R, Kim JH, et al. Association of the Female Athlete Triad Risk Assessment Stratification to the Development of Bone Stress Injuries in Collegiate Athletes. Am J Sports Med. 2017;45: 302–310. doi:10.1177/0363546516676262/ASSET/IMAGES/LARGE/10.1177_0363546516676262-FIG1.JPEG

47. Aladashvili-Chikvaidze N, Kristesashvili J, Gegechkori M. Types of reproductive disorders in underweight and overweight young females and correlations of respective hormonal changes with BMI. Iran J Reprod Med. 2015;13: 135. Available: /pmc/articles/PMC4426152/

48. Kraus E, Tenforde AS, Nattiv A, Sainani KL, Kussman A, Deakins-Roche M, et al. Bone stress injuries in male distance runners: higher modified Female Athlete Triad Cumulative Risk Assessment scores predict increased rates of injury. Br J Sports Med. 2019;53: 237–242. doi:10.1136/BJSPORTS-2018-099861

49. Golubnitschaja O, Liskova A, Koklesova L, Samec M, Biringer K, Büsselberg D, et al. Caution, “normal” BMI: health risks associated with potentially masked individual underweight—EPMA Position Paper 2021. EPMA J 2021 123. 2021;12: 243–264. doi:10.1007/S13167-021-00251-4

50. Cavanagh P, Williams K. The effect of stride length variation on oxygen uptake during distance running. Med Sci Sport Exerc. 1982;14: 30–35.

51. de Ruiter CJ, van Daal S, van Dieën JH. Individual optimal step frequency during outdoor running. https://doi.org/101080/1746139120191626911. 2019;20: 182–190. doi:10.1080/17461391.2019.1626911

52. Van Oeveren BT, De Ruiter CJ, Beek PJ, Van Dieën JH. Optimal stride frequencies in running at different speeds. PLoS One. 2017;12: e0184273. doi:10.1371/JOURNAL.PONE.0184273

53. Schiff MA, O’halloran R, Schiff MA, Caine DJ. Injury Prevention in Sports. http://dx.doi.org/101177/1559827609348446. 2009;4: 42–64. doi:10.1177/1559827609348446

54. Delp SL, Loan JP, Hoy MG, Zajac FE, Topp EL, Rosen JM. An interactive graphics-based model of the lower extremity to study orthopaedic surgical procedures. IEEE Trans Biomed Eng. 1990;37: 757–767. doi:10.1109/10.102791

55. Yamaguchi GT, Zajac FE. A planar model of the knee joint to characterize the knee extensor mechanism. J Biomech. 1989;22: 1–10. doi:10.1016/0021-9290(89)90179-6

56. Anderson FC, Pandy MG. A Dynamic Optimization Solution for Vertical Jumping in Three Dimensions. http://dx.doi.org/101080/10255849908907988. 1999;2: 201–231. doi:10.1080/10255849908907988

57. Anderson FC, Pandy MG. Dynamic optimization of human walking. J Biomech Eng. 2001;123: 381–390. doi:10.1115/1.1392310

58. Friederich JA, Brand RA. Muscle fiber architecture in the human lower limb. J Biomech. 1990;23: 91–95. doi:10.1016/0021-9290(90)90373-B

59. Wickiewicz, Wickiewicz TL. Muscle architecture of the human lower limb. Clin Orthop Relat Res. 1983;179.

60. Gait 2392 and 2354 Models - OpenSim Documentation - Global Site. [cited 28 Jul 2023]. Available: https://simtkconfluence.stanford.edu:8443/display/OpenSim/Gait+2392+and+2354+Models

61. Catelli DS, Wesseling M, Jonkers I, Lamontagne M. A musculoskeletal model customized for squatting task. Comput Methods Biomech Biomed Engin. 2019;22: 21–24. doi:10.1080/10255842.2018.1523396/SUPPL_FILE/GCMB_A_1523396_SM7370.DOCX

62. Lin YC, Pandy MG. Predictive Simulations of Human Sprinting: Effects of Muscle-Tendon Properties on Sprint Performance. Med Sci Sports Exerc. 2022;54: 1961–1972. doi:10.1249/MSS.0000000000002978

63. dit Falisse AM, Groote F De, Jonkers I. A Framework for Personalized Modeling and Predictive Simulation to Study Gait in Children with Cerebral Palsy. 2019.

64. Groote F De, Kinney AL, Rao A V, Fregly BJ. Evaluation of Direct Collocation Optimal Control Problem Formulations for Solving the Muscle Redundancy Problem. Ann Biomed Eng. 2016;44: 2922–2936. doi:10.1007/s10439-016-1591-9

65. Bhargava LJ, Pandy MG, Anderson FC. A phenomenological model for estimating metabolic energy consumption in muscle contraction. J Biomech. 2004;37: 81–88. doi:10.1016/S0021-9290(03)00239-2

66. Kingma DP, Ba JL. Adam: A Method for Stochastic Optimization. 3rd Int Conf Learn Represent ICLR 2015 - Conf Track Proc. 2014 [cited 6 Jul 2023]. Available: https://arxiv.org/abs/1412.6980v9

67. Falisse A, Serrancolí G, Dembia CL, Gillis J, Groote F De. Algorithmic differentiation improves the computational efficiency of OpenSim-based trajectory optimization of human movement. Srinivasan M, editor. PLoS One. 2019;14: e0217730. doi:10.1371/journal.pone.0217730

68. Andersson JAE, Gillis J, Horn G, Rawlings JB, Diehl M. CasADi: a software framework for nonlinear optimization and optimal control. Math Program Comput. 2019;11: 1–36. doi:10.1007/s12532-018-0139-4

